# Classifying protein structures into folds by convolutional neural networks, distance maps, and persistent homology

**DOI:** 10.1101/2020.04.15.042739

**Authors:** Yechan Hong, Yongyu Deng, Haofan Cui, Jan Segert, Jianlin Cheng

## Abstract

The fold classification of a protein reveals valuable information about its shape and function. It is important to find a mapping between protein structures and their folds. There are numerous machine learning techniques to predict protein folds from 1-dimensional (1D) protein sequences, but there are few machine learning methods to directly class protein 3D (tertiary) structures into predefined folds (e.g. folds defined in the SCOP database). We develop a 2D-convolutional neural network to classify any protein structure into one of 1232 folds. We extract two classes of input features for each protein: residue-residue distance matrix and persistent homology images derived from 3D protein structures. Due to restrictions in computing resources, we sample every other point in the carbon alpha chain to generate a reduced distance map representation. We find that it does not lead to significant loss in accuracy. Using the distance matrix, we achieve an accuracy of 95.2% on the SCOP dataset. With persistence homology images of 100 × 100 resolution, we achieve an accuracy of 56% on SCOPe 2.07 dataset. Combining the two kinds of features further improves classification accuracy. The source code of our method (PRO3DCNN) is available at https://github.com/jianlin-cheng/PRO3DCNN.

## 1 Introduction

Structural Classification of Proteins (SCOP) is a database of protein structural relationships. The proteins are placed in a fold classification hierarchy in relative to their structural and evolutionary relations [1]. SCOP was initially a manually curated ordering of proteins [1]. However, as numerous proteins continue to be discovered at a rapid pace, a need for an automatic method of classification became necessary. Structural Classification of Proteins – extended (SCOPe), extends the original SCOP database by using some structure/sequence comparison methods to classify newly discovered proteins [1].

Similar to other complex taxonomies, SCOP classification scheme was created as a hierarchy with a basis in evolutionary relationships. SCOP characterizes each of these levels in the hierarchy with the following description: *family* containing proteins with similar sequences and close evolutionary relationships; *superfamily* bridging together protein families with common functional and structural features inferred to be from a common evolutionary ancestor; *folds* grouping structurally similar superfamilies that may not have evolutionary relationships, and *classes* based mainly on secondary structure content and organization in protein tertiary structures. The highest level of the hierarchy, class, consists of 7 categories. Alpha proteins (a), beta proteins (b), Alpha/Beta proteins (c), Alpha+Beta proteins (d), multi-domain proteins (e), membrane and cell surface proteins and peptides (f), and small proteins (g) [2].

Many works have been done on using the protein sequences and their evolutionary relationships with each other to predict structural folds of proteins. This is primarily due to the lack of structural information for many newly discovered proteins. Sequence based classification using deep convolutional neural networks like DeepSF show that it is possible to predict protein fold with an accuracy up to 75.3% [3].

In this paper, we shift the focus from the protein’s sequence to the topological structure to predict the protein’s fold. Since each fold belongs to only one class, predicting the protein fold also predicts the protein’s class. We introduce two methods for classifying protein fold. First, we use a residueresidue distance matrix as an input matrix to a convolutional neural network. Second, we use mathematical persistent homology (note: a mathematical concept different from sequence homology in the bioinformatics field) to extract topological features of protein structures and plot these features as an input matrix to a convolutional neural network. The background of the two kinds of features and related work are described below.

### Distance Matrix

Each protein has a backbone structure that is formed by series of connecting points. The backbone structure of the protein is characteristic of the protein’s general shape. A distance matrix of the protein’s backbone is created, where the r^th^ row and c^th^ column of the matrix gives the distance between the r^th^ point and the c^th^ point of the protein backbone. Then this matrix is used as an input to our convolutional neural network. We take a look at few of the representative proteins from alpha proteins, beta proteins, alpha/beta proteins, and alpha+beta proteins. We juxtapose each protein’s distance matrix to the protein’s structure to see if there are significant differences between the classes (**Figure 1**). It is shown that the distance matrices do represent the alpha helices, the beta sheets, and also the relationship between these features.

**Figure 1.**
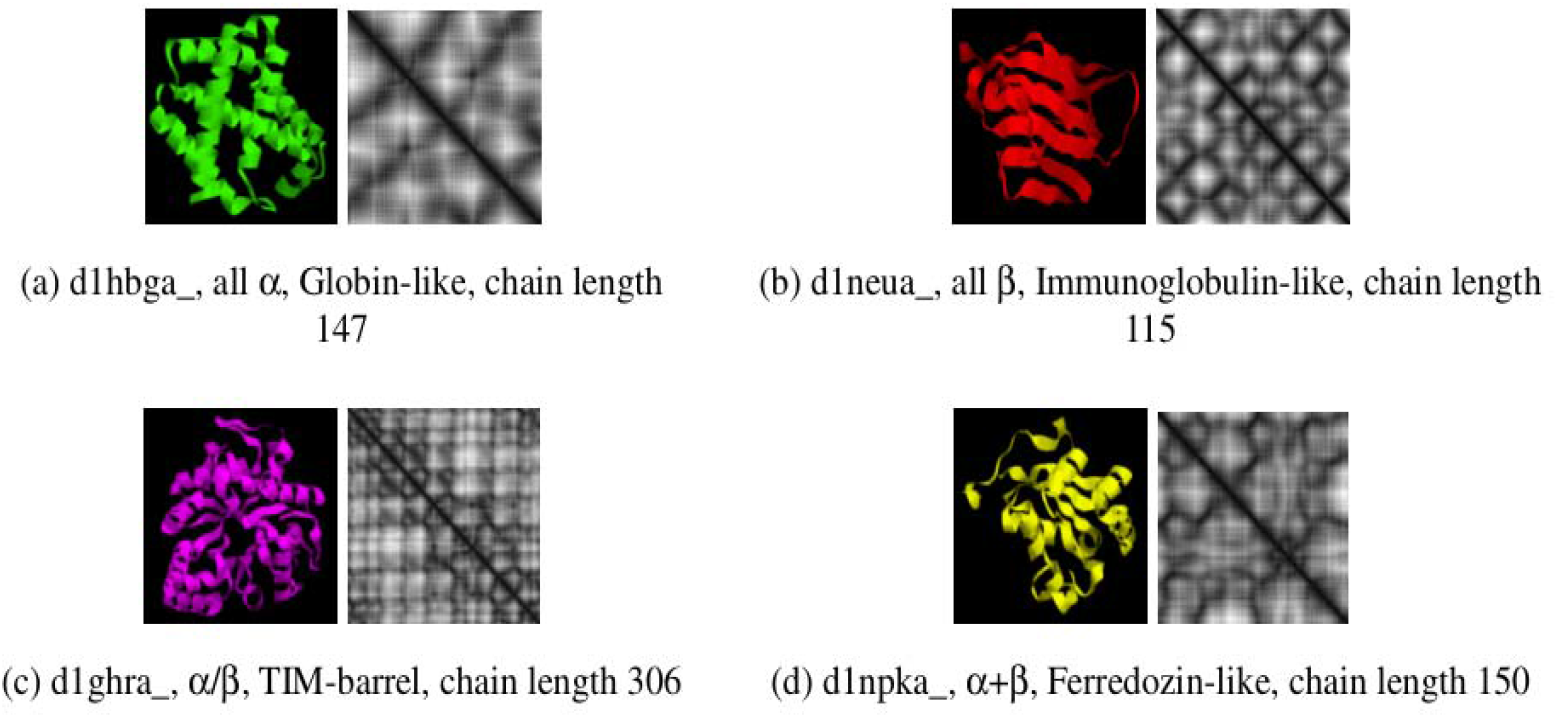
A comparison of the distance matrices of several proteins of different classes from the SCOP database. We see that the distance matrix is able to pick up on characteristic alpha helix and beta sheets of each protein, making it likely that this feature will perform well.

There has been existing work (Fast SCOP Classification) on using the distance matrix to classify proteins from a very small subset of SCOP database [4] of 698 proteins. The distance matrix is computed for the protein and fed into a multi-class support vector machine (SVM) of the One-Versus-One (OVO) variant. This approach achieved an accuracy of 74.55% on this small subset of SCOP.

Fast SCOP Classification extracts regions of interest (ROI) from the distance matrix by decomposing the upper triangle of the distance matrix into ROI (**Figure 2**). These regions attempt to capture the regions of the distance matrix where there are alpha helices and beta sheets. The relationships between these regions of interest are abstracted into a feature set, which is given to the SVM for fold classification.

**Figure 2.**
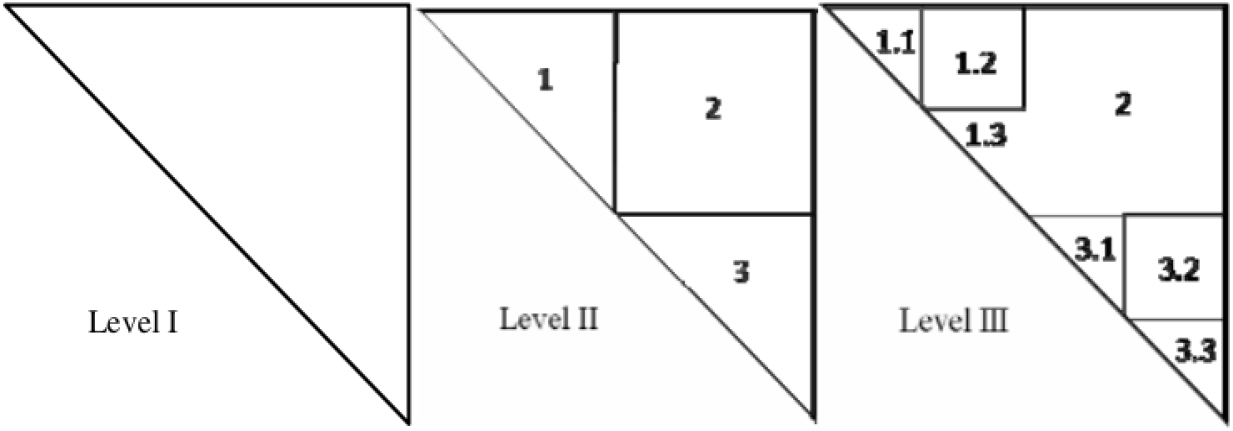
Fast SCOP Classification decomposing the upper triangle of a distance matrix into regions of interest (ROI) to extract important features such as alpha helices and beta sheets.

In comparison, we employ a convolutional neural network to have the model learn the ROIs on it’s own. The CNN has much more advantages over the Fast SCOP’s method because complex relationships between regions of interest can be modeled by the hidden layers automatically. Also the regions of interests are dynamically formed as the model is trained. There are no supervisions into constructing the high-level feature set of ROIs, allowing the model to learn complex behaviors that may be elusive or difficult to define clearly. However, a downside to the CNN is that it is more computationally intensive, requiring a long training time and a large set of data to learn effectively.

### Persistent Homology

Persistent homology is a mathematical theory from algebraic topology that allows the characterization of a point cloud by the observing creations of holes, which we call cycles, in the data and how long these cycles persist. We iterate over a real number variable, ε, and connect the edges in our point cloud whose length is smaller than ε. A hole is formed when a closed loop is formed by the path of the edges. In the example with 4 points (**Figure 3**), a cycle is formed when the four edges of the rectangle’s boundary is formed. The ε value at which the cycle is formed is called the birth of this cycle. This cycle is destroyed when triangles formed within our cycle to completely fill it in. The ε value associated with the cycle being filled in is called the death of the cycle. We say that the cycle persists for a duration of birth – death. Each cycle forms an interval, [birth, death]. The collection of all intervals from the cycles in the data is called a barcode.

**Figure 3.**
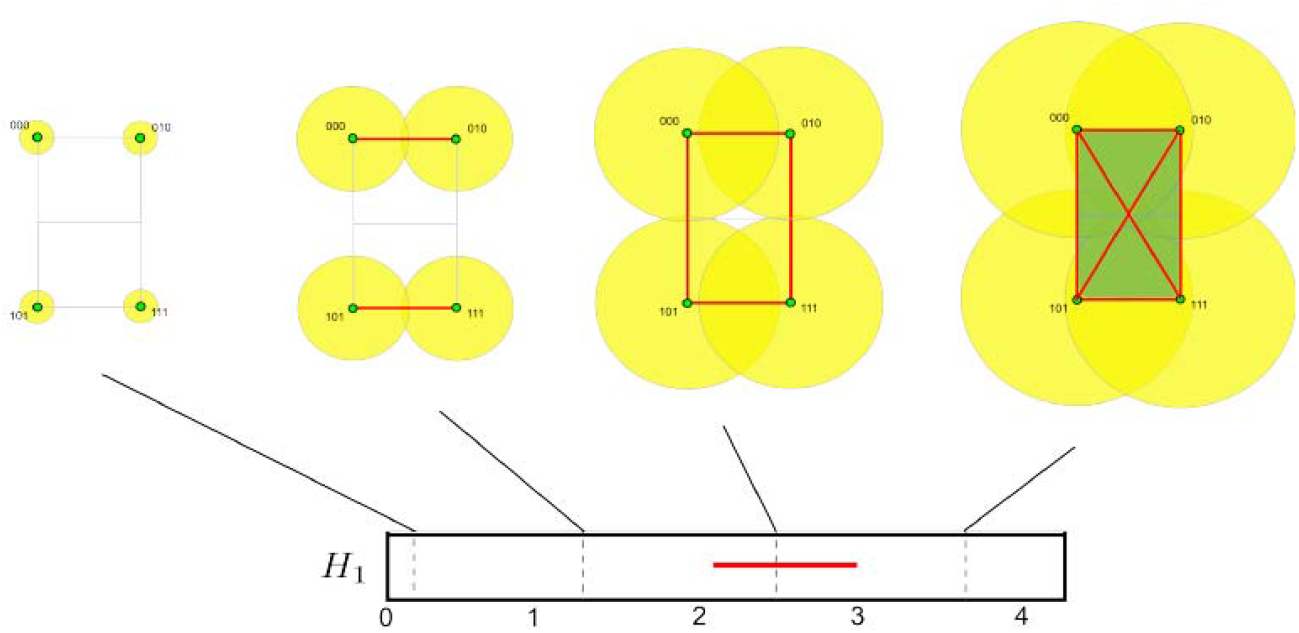
The persistent homology characterizing the point cloud of four points. A rectangular cycle is formed when the 4 boundary edges of the rectangles are formed at ε = 2. We see that this rectangular cycle is dies at ε = 3

When we observe the barcode of the proteins from the different classes, we do see some distinguishing patterns (**Figure 4**). The proteins with alpha helices have a long-curved edge towards the top of the barcode, while proteins with beta sheets have a thicker bar towards the bottom.

**Figure 4.**
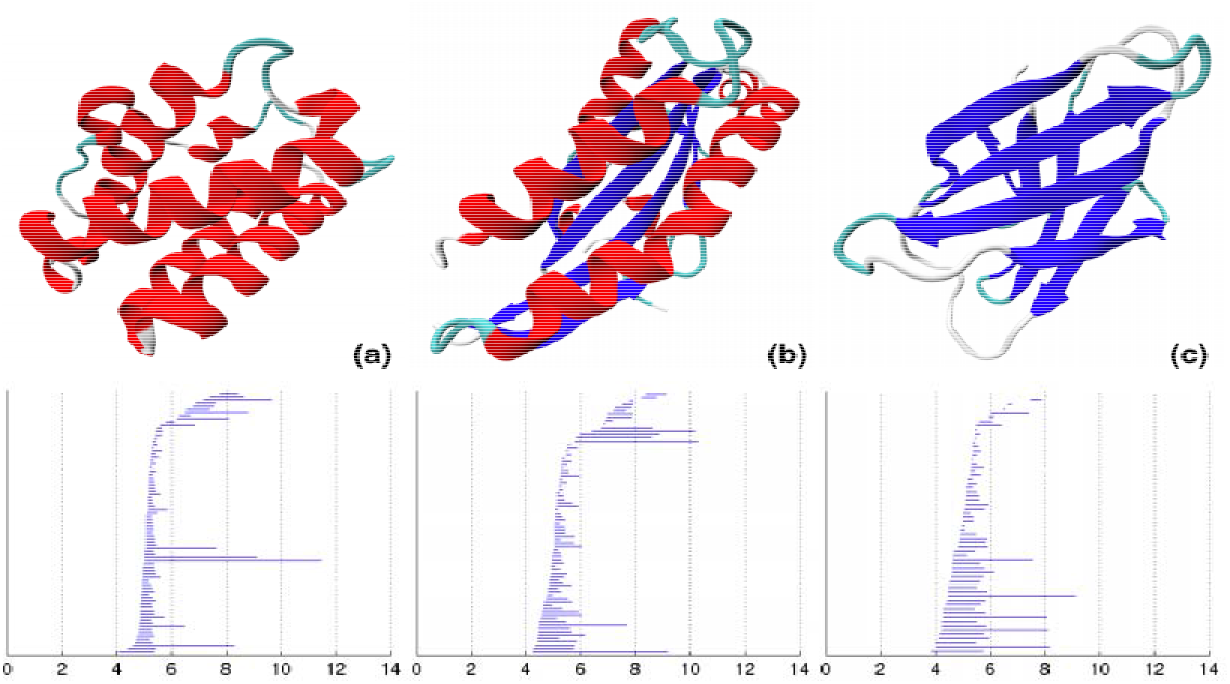
Persistent homology was used to calculate the barcodes on three proteins from Alpha protein (a), Alpha/Beta protein (b), Beta protein(c) classes. The proteins with alpha helices (red) have elongated structures at the top of the barcode while proteins with beta sheets (blue) are thick at the base.

There have been existing attempts at using persistent homology to classify protein classes, the highest level of SCOP hierarchy [2]. Cang et al. [2] applies persistent homology on a small set of 900 proteins selected from Alpha, Beta, and Alpha/Beta classes. Each class is represented by 300 proteins and 60 proteins were selected for testing. An SVM is used to classify the classes with an average accuracy of 85%.

In comparison, we classify proteins in to a large number of folds, a more specific level of the SCOP hierarchy. Our task of classification is much more extensive than the classification task performed in Cang. We classify 276,231 proteins into 1232 labels compared to the 900 proteins into 3 labels. We also use a convolutional neural network as opposed to a SVM.

Based on our experiments of using the two kinds of features, we note that the distance matrices contain much more information than the barcodes. This difference in resolution and details may cause the distance matrices to perform better than the barcodes.

## 2 Materials

The training, validation and test data are generated from SCOP and SCOPe database, the databases of proteins organized into hierarchical classes based on their shape and function. There are 4 levels of the important hierarchy (top down): Class, Folds, Superfamily, Family. We will be primarily concerned with the Fold.

SCOPe 2.07 is a database of 276,231 proteins that have been organized into 7 Classes, 1232 Folds, 2026 Superfamilies, and 4919 Families. The dataset was released and updated till 2017. This dataset contains and is about 8 times larger than the SCOP 1.55 dataset. The statistics on the number of classes, folds, superfamilies, and families is shown in **Table 1**. **Figure 5** is a histogram showing the distribution of the number of proteins in each fold. **Figure 6** illustrates the distribution of the protein lengths. We split up 70% of the dataset for training, 15% for validation and 15% for testing. We adjust the sampling of the validation and testing so that a wide range of folds are represented.

**Table 1.** The statistics of 276231 entries of SCOPe 2.07.

| Class | Number of folds | Number of superfamilies | Number of families |
| --- | --- | --- | --- |
| a. Alpha proteins | 289 | 516 | 1062 |
| b. Beta proteins | 178 | 370 | 968 |
| c. Alpha and Beta proteins (a/b) | 148 | 246 | 986 |
| d. Alpha and beta proteins (a+b) | 388 | 565 | 1338 |
| e. Multi-domain proteins (alpha and beta) | 71 | 71 | 118 |
| f. Membrane and cell surface proteins and peptides | 60 | 119 | 173 |
| g: Small proteins | 98 | 139 | 274 |
| **Total** | **1232** | **2026** | **4919** |

**Figure 5.**
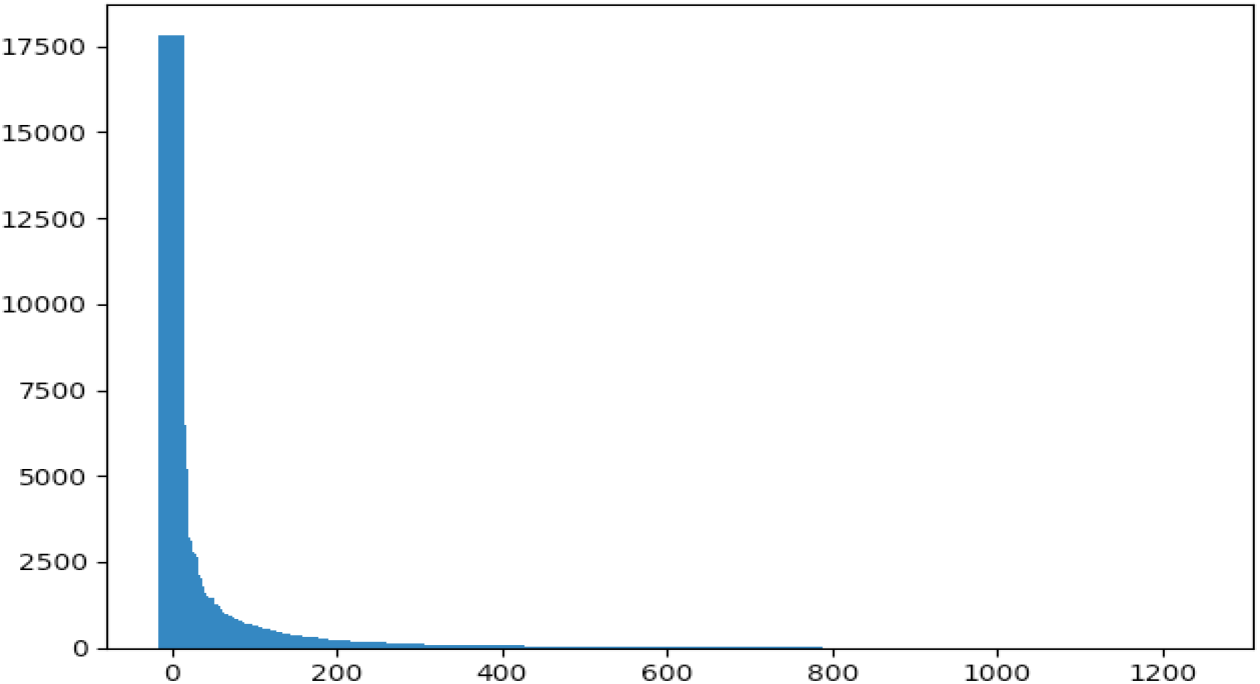
The histogram of the number of proteins per fold for SCOP 2.07. x-axis is the index of folds and y-axis the number of proteins in each fold.

**Figure 6.**
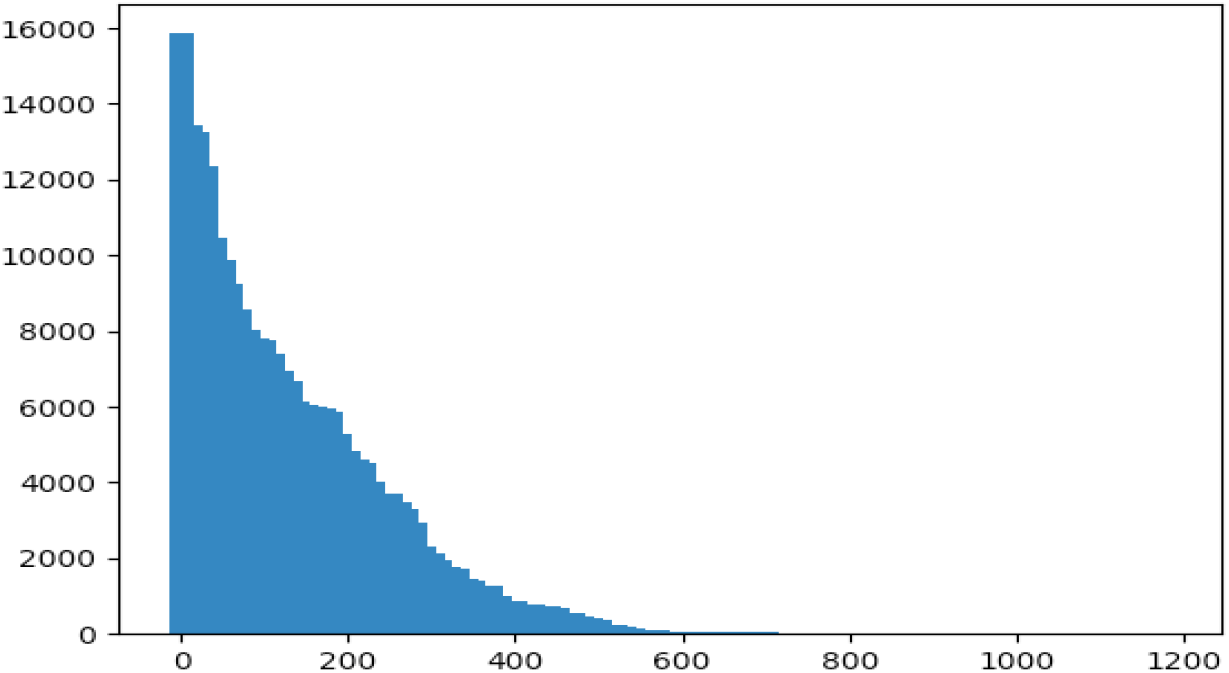
The histogram of the protein lengths for SCOP 2.07. x-axis is the length of proteins and y-axis the number of proteins at each length. We see that 90% of the proteins have length less than 500.

## 3 Methods

### 3.1 Generation and Analysis of Distance Matrix

With the points in the protein backbone, we construct a distance matrix of the distances between the points. The unit of distance provided in the protein PDB fi ms. We used the Euclidean distance for the distances between the points.

We analyze the distance matrix to see if important structural features are represented in the matrix. To help with visualization, the distance matrix is mapped to an image of equal size, where closest distances appear in white and furthest distances appear in black (**Figure 7**). Distances in between take a gray hue with the intensity based on its value. Along the diagonal line of the image, the distance matrix is completely white. This is because the distance between a point to itself is zero. We note that this diagonal white line uniquely identifies the protein backbone chain (since only the distance between a point and itself has the distance of zero). Having a clear representation of the protein backbone is important because it is a central structure that other features can be spatially oriented around.

**Figure 7.**
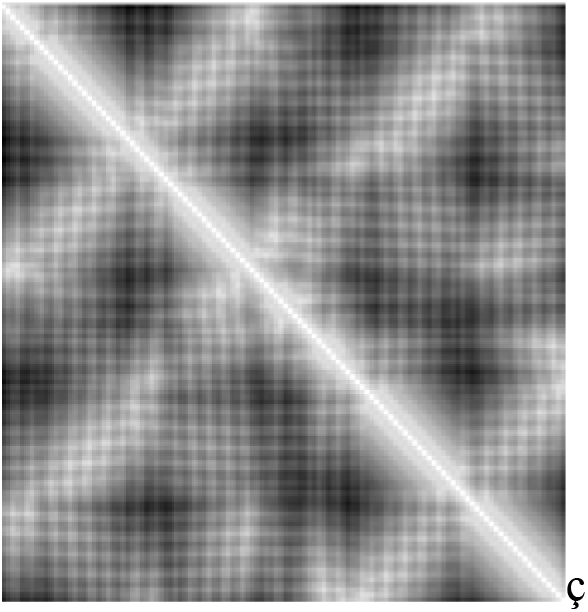
The distance matrix of the protein 1ux8. We see a thick diagonal line with corresponds to the protein backbone chain.

We note thick white regions running parallel along the diagonal line of the image (**Figure 8**). These regions indicate that at a given point in the chain, it is in close proximity to the nearby neighbors. We also note that in for a point in an alpha helix, it is also in close proximity to its nearby neighbors. In our example, these four thick white regions correspond the four helix structures on our protein.

**Figure 8.**
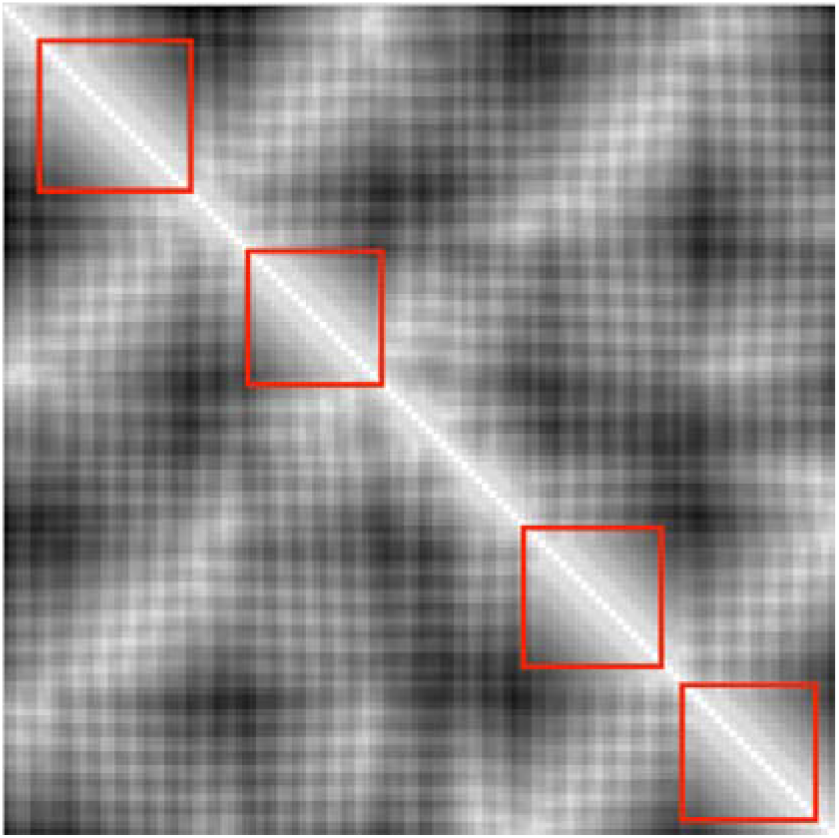
The red squares outline the discrete thick white regions running parallel to the diagonal line. These regions indicate that the neighbors of points in this region are in very close proximity, indicating that there may be a helix.

We note patches of thick white lines in the intersection of the rows belonging to a beta strand and the columns belonging to another beta strand (**Figure 9**). This indicates that the points of the two strands are in close proximity. In particular, the regions are close sequentially: the i^th^ point in strand A is close to the j^th^ point in strand B and the i+1^th^ point in strand A is close to the j-1^th^ point in strand B. This sequential relationship describes an anti-parallel beta sheets (perpendicular to the diagonal). For parallel beta sheets, the i+1^th^ point would be close to the j+1^th^ point (in parallel to the diagonal).

**Figure 9.**
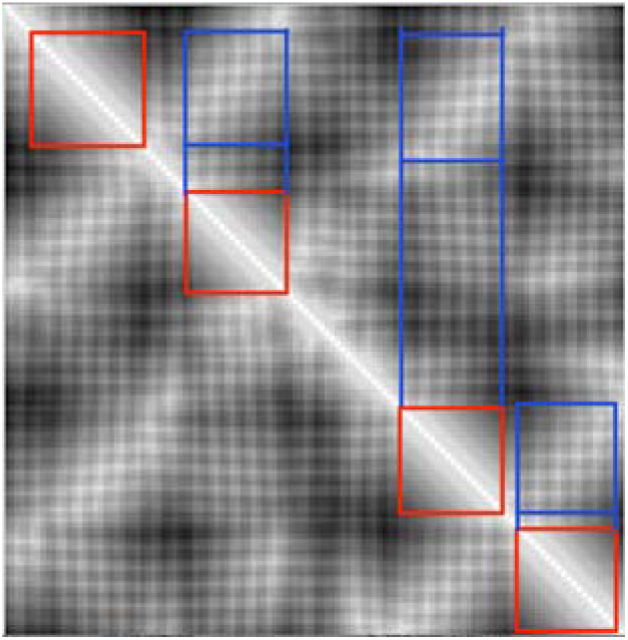
The blue squares outline the regions of proximity between points on different regions, suggesting that the interaction of beta-strands. The direction of the lines indicates whether the two strands are parallel or anti-parallel.

### 3.2 Generation of Persistent Homology for Protein Structures

We study Persistent Homology of the set of points in a protein backbone. Persistent Homology produces a barcode invariant, or alternately by a persistence image, that encodes the significant topological features of the data. Topological Data Analysis uses persistent homology to study point cloud data. A point cloud is a set of points in Euclidean space. We are interested in the point cloud consisting of protein backbone points in 3-dimensional space

The barcode invariants of persistent homology faithfully encode essential features of a point cloud. Persistence images retain these important features, and therefore make excellent feature vectors for machine learning with point cloud data. First, barcodes are invariant under rotation and translation of the entire point cloud. Second, barcodes are “stable” in a specific technical sense under small perturbations, or measurement errors, of the positions of the individual points comprising a point cloud. In particular, the intervals of greater length (persistence) are insensitive to perturbation, and represent significant features of the point cloud. Conversely the intervals of short length represent less important ephemeral features, which may appear or disappear under perturbation.

We apply persistence homology to our data set. Here we use a protein 1ux8 as an example to illustrate the idea. We generate the barcode for a protein 1ux8 and see if the backbone chain, alpha helices, and beta sheets are represented in the barcodes. We also compare the features generated by persistence homology and the distance matrix. We inspect holes generated by persistence homology of protein 1ux8, a protein of length 118. At each point on the protein backbone, we search an ε-radius around the point to connect edges of increasing length to form the ε-Vietoris-Rips complexes. **Figures 10, 11**, and **12** shows a series of ε-Vietoris-Rips complexes.

**Figure 10.**
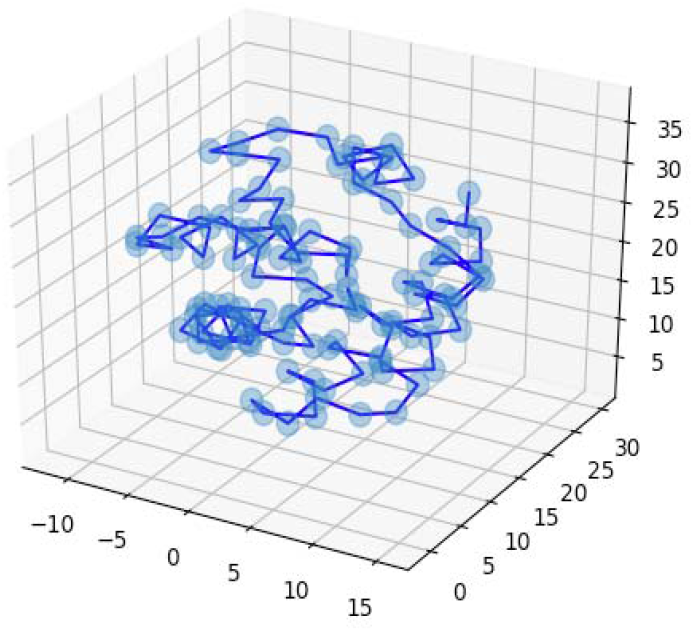
Atε = 1, a search is done at each point at increasing radii to connect the nearby points

**Figure 11.**
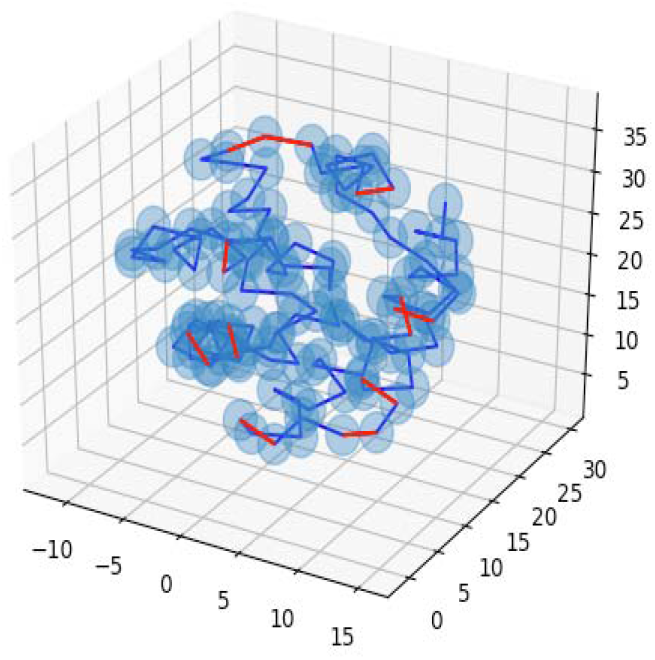
Atε = 3.5, the red edges indicate that the two points are less than 3.5 distance away, making them connected. As we increase the ε, we connect more and more points, forming more simplexes.

**Figure 12.**
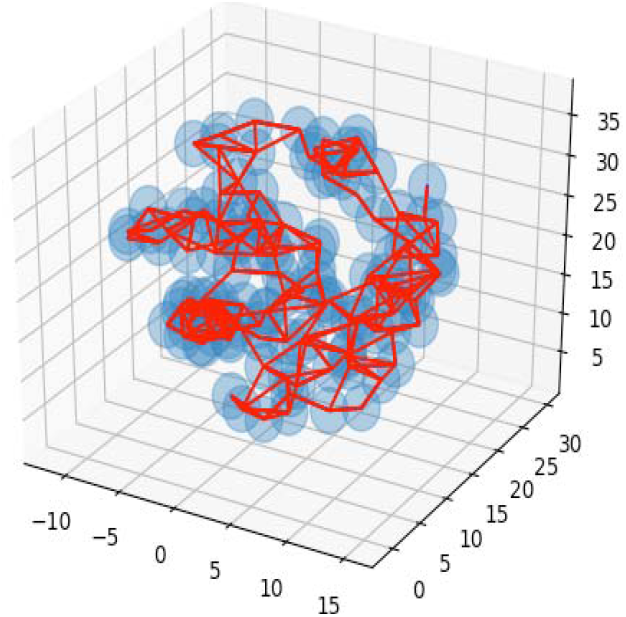
The general structure of the protein is detected as we increase when ε.

After a certain point we want to cut off our search radius because we have extracted all the notable

We inspect some key holes generated by the persistent homology to see if it captures the features of our data well. We see that persistent homology is able to detect the presence of alpha helices (e.g. **Figure 13**) and also beta sheets. The persistent homology is almost on par with the distance in terms of representing alpha helices and beta sheets. However, the information of the birth and the death of the holes alone is not enough to describe spatial relationships between multiple pairs of alpha helices.

**Figure 13.**
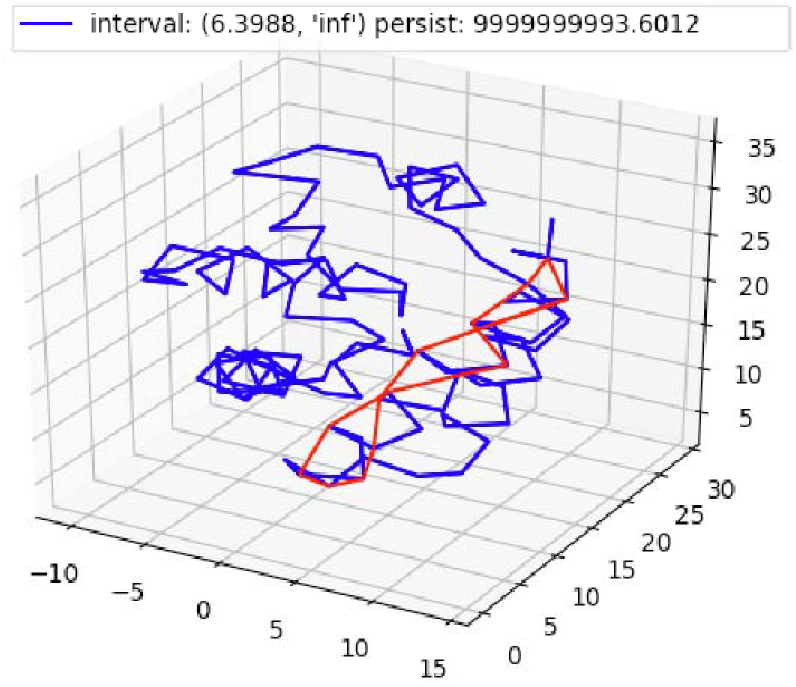
Persistent homology detecting an alpha helix. The hole is a significant feature of our data since the hole persists for a long time.

The computation of persistent homologies for 1/10 protein structures of SCOP 1.55, took around 15 hours. Upon inspection of the computation times, we noted that proteins that took longer than 50 seconds to compute accounted for about 11 hours of the 15 hours. The reason why persistent homology had a difficult time was due to an improperly set cutoff value. Initially we set the cutoff value as the maximum length of the edges found in the protein backbone. This was because we wanted to include the edges of the protein backbone as part of the filtration of ε Vietoris-Rips simplicial complexes because it is a very important feature of the protein. However, some of the proteins had very long edges in the protein backbone chain, making the cutoff value very large and making our algorithm compute the persistent homology for all the edges in our protein. When we adjusted the cutoff value to be just around, we noticed that beta helices along the protein were not being captured very well by the holes. To remedy this issue, we multiply a weight of .25 on the edges belonging to the protein backbone. This would ensure that protein backbone edges that are reasonably small would get included as part of the filtration of ε Vietoris-Rips simplicial complexes. Placing this modification greatly sped up the speed of the computation of persistent homologies, computing 1/10 of SCOP 1.55 from 15 hours to 2.6 seconds.

### 3.3 Convolutional Neural Network (CNN) Model

**Figure 14** illustrates the CNN architecture to classify protein structures into folds using distance matrices, persistence homology or both as input. It is a relatively simple network. However, for our task of classifying protein structures, it performs exceptionally well. Because of its simple architecture, it less likely overfits the data and can be trained at a fast speed. For all of the training, a batch size of 50 was used. Stochastic gradient descent with a learning rate of .01 was used. The labels of the protein folds were one-hot encoded as a vector of size 605 for SCOP 1.55 and 1232 for SCOPe 2.07. For SCOP 1.55 the models were trained for 20 epochs. SCOPe 2.07, the models were trained for 30 epochs.

**Figure 14.**
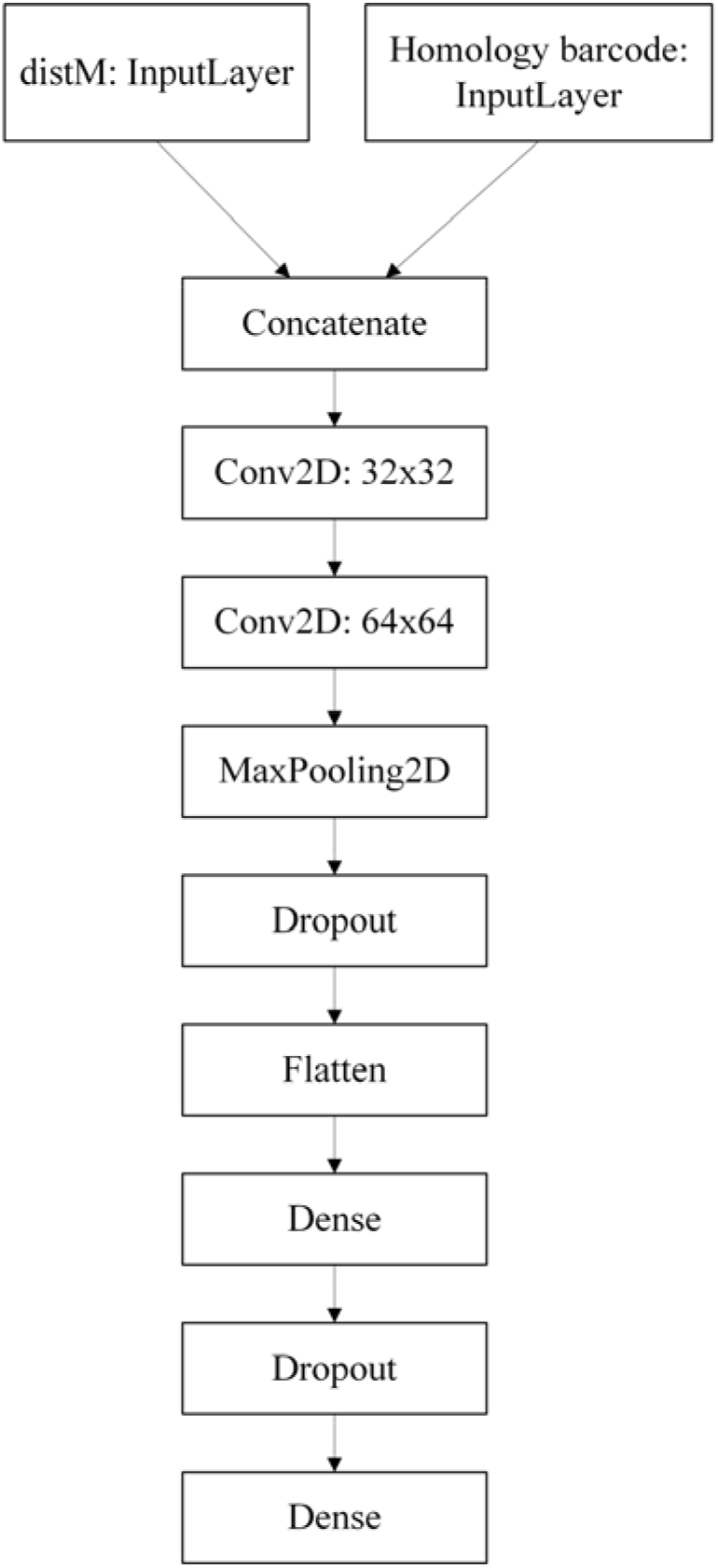
The convolutional neural network for classifying protein structural information represented by distance matrix, persistency homology, or both into folds.

During the experiment, the accuracy is higher using distance matrix than using persistence homology. It is probably because persistence homology captures less topological features than distance matrices. To further optimize the accuracy and robustness of the model, we experimented the combination of distance matrix and persistence homology as the input of our convolutional neural network. The persistence images are still set to 100×100 resolution. Since we are using a 100×100 window to crop our distance matrix, some long proteins will have more than 1 input matrix. To solve the asymmetry of input features, we duplicate the corresponding amount of persistence images. For example, to train a protein with a length of 600, we crop the distance matrix to 4 100×100 matrices and duplicate 4 copies of 100×100 persistence image.

## 4 Results and Discussions

### 4.1 Training and validation on SCOP 1.55 and SCOPe 2.07 dataset

We trained, validated and tested the deep convolutional neural network on SCOP 1.55 and SCOPe 2.07 dataset. The results on the test dataset are reported in **Table 2**. Overall, the distance matrix outperforms persistent homology on both SCOP 1.55 and SCOPe 2.07. On SCOP 1.55 the distance matrix has an accuracy of 87% and the persistence homology has an accuracy of 62%. We also tried both distance matrix and persistent homology together as a combined input feature, and the combined input has an accuracy of 91% on SCOP 1.5 dataset, which is higher than the individual inputs. On SCOPe 2.07, the distance matrix has an accuracy of 95.2 % and the persistence homology has an accuracy of 56%. Combining the two also slightly improves the prediction accuracy.

**Table 2.** The classification accuracy of using different inputs on SCOPe 2.07 and SCOP 1.55 datasets

| Method | Accuracy on SCOPe 2.07 | Accuracy on SCOP 1.55 |
| --- | --- | --- |
| Distance Matrix | 95.2% | 87% |
| Persistence Homology | 56% | 62% |
| Combined Input | 95.6% | 91% |

The distance matrix performs a lot better than the persistence homology because the distance matrix conveys the spatial relationship between the alpha helices and beta sheets while the persistence images only identify their existence. Specifically, persistence homology would have trouble distinguishing between proteins of Alpha/Beta and Alpha+Beta classes since these proteins are both composed of alpha helices and beta sheets but their spatial orientations are different.

We notice that the number proteins in a fold influence the classification accuracy. **Figure 15** plots the classification accuracy for a fold against the number of proteins in the fold. There is a positive correlation between the two.

**Figure 15.**
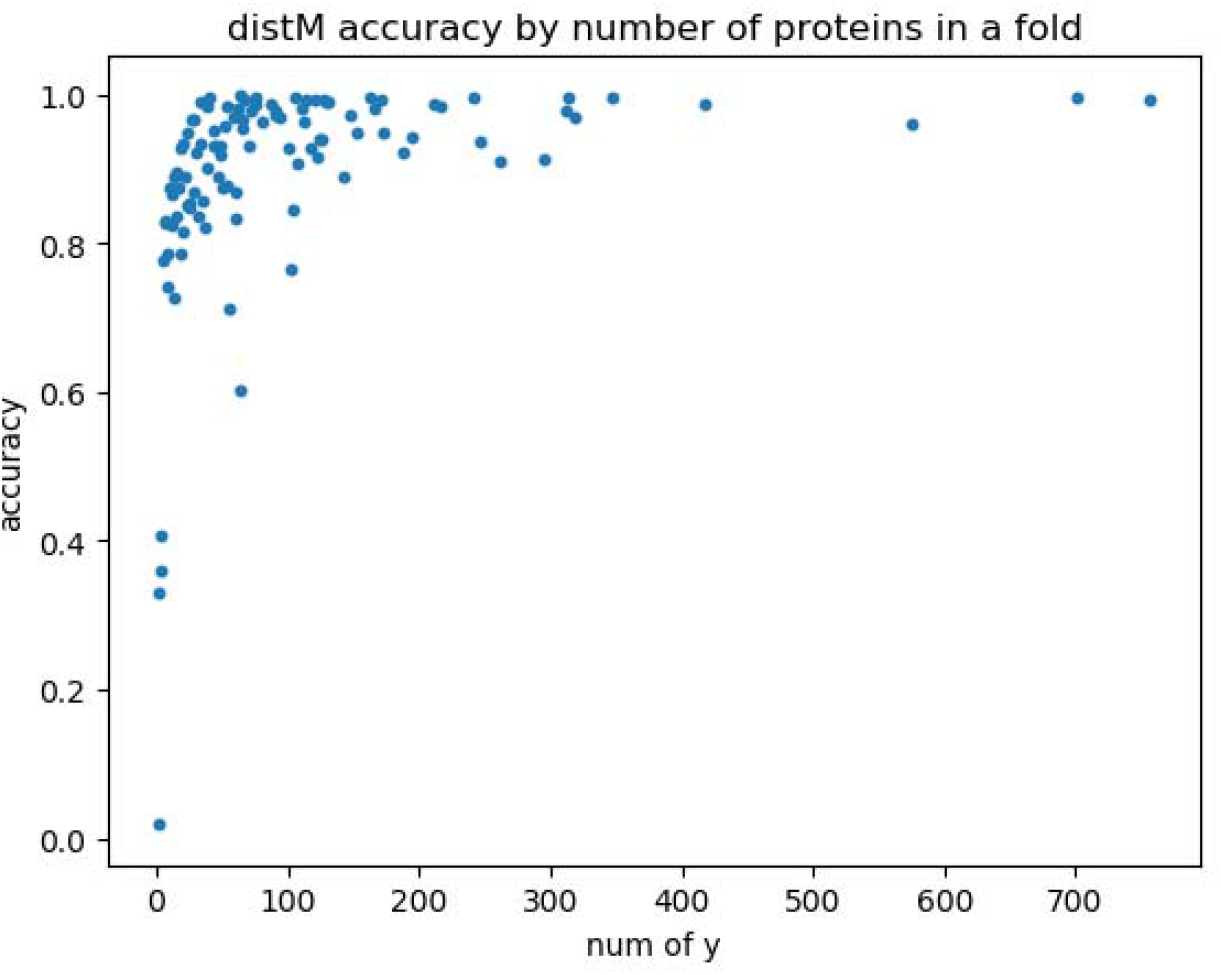
A plot of classification of accuracy against the number of proteins in a fold using the distance matrix input on SCOP 1.55 dataset.

The persistence homology performs worse in SCOPe 2.07 than SCOP 1.55. Although there are more training data per fold in SCOPe 2.07, there are more proteins that require more information about the spatial orientation of beta sheets and alpha helices that were added in SCOPe 2.07. This may make it more difficult for persistence homology to be accurate.

We found that the sparse distance matrix reduced by two folds and the regular distance matrix perform similarly. The sparse distance matrix does take more epochs during training for it to reach the similar level of accuracy as the regular distance matrix.

## 5 Conclusion

We presented a deep convolution neural network to classify a protein topological feature into one of all 1232 folds defined in SCOPe 2.07 using distance matrices and persistence homology generated from protein structures. To our knowledge, this is the first system that can directly classify proteins from the topological space to the entire fold space rather accurately without using sequence or structure comparison. Using distance matrix and persistence homology, our method is not only complementary with traditional sequence-alignment methods based on pairwise comparison, but also provides a new way to study protein structures.

